# Linking Altered Neuronal and Synaptic Properties to Nicotinic Receptor Alpha5 Subunit Gene Dysfunction: A Translational Investigation in Rat mPFC and Human Cortical Layer 6

**DOI:** 10.1101/2024.04.12.589167

**Authors:** Danqing Yang, Guanxiao Qi, Daniel Delev, Uwe Maskos, Dirk Feldmeyer

## Abstract

**Background:** Genetic variation in the α5 nicotinic acetylcholine receptor (nAChR) subunit of mice results in behavioral deficits linked to the prefrontal cortex (PFC). A Single Nucleotide Polymorphisms (SNP) in CHRNA5 imparts a partial loss of function to the α5 subunit-containing (α5*) nAChRs and have been demonstrated to be associated with psychiatric disorders in humans, including schizophrenia, nicotine dependence, cocaine and alcohol addiction.

**Methods:** We performed single cell-electrophysiology combined with morphological reconstructions on layer 6 (L6) excitatory neurons in the medial PFC (mPFC) of wild type (WT) rats (n = 25), rats carrying the human coding polymorphism rs16969968 in *Chrna5* (n = 11) and α5 knockout (KO) rats (n = 28). Neuronal and synaptic properties were compared among three rat genotypes. Galantamine was applied to identified L6 neuron populations to specifically boost the nicotinic responses mediated by α5*nAChRs in the rat mPFC and human neocortex (n = 6 patients).

**Results:** Compared with neurons in WT rats, L6 regular spiking (RS) neurons in the α5KO group exhibited altered electrophysiological properties, while those in α5SNP rats remained unchanged. L6 RS neurons in mPFC of α5SNP and α5KO rats differed from WT rats in dendritic morphology, spine density and spontaneous synaptic activity. Galantamine acted as a positive allosteric modulator of α5*nAChRs in RS but not burst spiking (BS) neurons in both rat and human cortical L6.

**Conclusion:** Our findings suggest that dysfunction in the α5 subunit gene leads to aberrant neuronal and synaptic properties, shedding light on the underlying mechanisms of cognitive deficits observed in human populations carrying α5SNPs. They highlight a potential pharmacological target for restoring the relevant behavioral output.

## Introduction

Acetylcholine (ACh) is believed to play a critical role in memory processing, arousal, attention, learning and sensory signaling through activating both muscarinic and nicotinic ACh receptors (AChRs) [1–5]. The medial prefrontal cortex (mPFC) is one of the various brain cortices affected by cholinergic modulation, and it is essential for working memory and top-down attention [6, 7]. The mPFC is densely innervated by cholinergic axon from the basal forebrain, and ACh release in this brain area has been linked to cue detection and attentional performance [8–10]. Schizophrenia (SZ) is characterized by impairment of cognitive processes that are critically dependent on the mPFC [11]. The mPFC is a crucial component of default mode network (DMN), which represents a neural baseline and is negatively correlated with task-positive activity [12]. Studies have reported reduced activity in the DMN and hypofrontality in SZ patients [13–15]. However, the cellular and circuit deficits underlying these phenomenon remain largely unexplored.

nAChRs are transmembrane ionotropic receptors activated by ACh and nicotine. They are central to nicotine addiction [16]. Previous studies have demonstrated a robust nAChR-induced response in pyramidal cells in deep neocortical layers [17–19]. These neurons prominently express hetero-pentameric α4β2* nAChRs, some of which contains the α5 subunit encoded by the *Chrna5* gene [20, 21]. The co-assembly of α5 subunits into α4β2 nAChRs enhances the Ca^2+^ permeability of the receptor, increases the receptor affinity for ACh, and slows down the receptor desensitization [22–24].

Several disease have been linked to coding and non-coding ***S***ingle ***N***ucleotide ***P***olymorphisms (***SNP***s) in CHRNA5 in a number of comprehensive ***G***enome-***w***ide ***A***ssociation ***S***tudies (***GWAS***) (for a review see [25]). In addition to a robust association with smoking [26], these SNPs are implicated in conditions such as SZ [27], addiction to cocaine [28], opioids [29], alcohol [30], cannabis [31], as well as chronic obstructive pulmonary disease (COPD) [32], lung cancer [33], longevity [34] and body-mass index [35]. The α5SNPs are highly prevalent in humans, with an average allele frequency of 28% in the general population, making them a major target for pharmacological intervention.

In mice, genetic deletion of the α5 nAChR subunit results in deficits in behavior controlled by the PFC, accompanied by altered cholinergic excitation and aberrant neuronal morphology in layer 6 (L6) of the PFC [24, 36–38]. However, it remains unclear whether α5SNP can induce neuronal pathophysiology and altered nicotinic signaling in this neuron population. In this study, we performed whole-cell recordings on L6 excitatory neurons in the rat mPFC using a transgenic rat line expressing a human α5SNP rs16969968 [39] and compared it with wild-type and *Chrna5* knock-out rats. This approach allowed us to study the effect of this variant on neuronal electrophysiology, morphology and nicotinic signaling. A positive allosteric modulator (PAM) specifically targeting α5*nAChRs in rat and human cortical L6 was used to investigate the mechanisms underlying the cognitive deficits observed in human α5SNPs, with the aim of identifying pharmacological effectors that could potentially restore behavioral performance.

## Methods

### Animals and patients

All experimental procedures involving animals were performed in accordance with the guidelines of the Federation of European Laboratory Animal Science Association, the EU Directive 2010/63/ EU, and the German animal welfare law. Wild type (WT) Long-Evans rats were purchased from Janvier Lab. α5KO and α5SNP Long-Evans rats were generated and housed at Institut Pasteur, Paris, France [39] and homozygous rats were bred at Research Centre Juelich, Juelich, Germany. Male and female Long-Evans WT (n = 25), α5KO (n = 28) and α5SNP (n = 11) rats aged 28-105 postnatal days were used for experiments. All rats were individually housed in a temperature-controlled environment on a 12-h reverse light-dark cycle, with food and water available ad libitum.

All patients underwent neurosurgical resections because of hippocampus sclerosis or tumor removal. Written informed consent to use spare neocortical tissue acquired during the surgical approach was obtained from all patients. The study was reviewed and approved by the local ethic committee (EK067/20). The cases were meticulously selected to fulfil two main criteria: 1) availability of spare tissue based on the needed surgical approach; and 2) normal appearance of the tissue according to radiological and intraoperative criteria (absence of edema, absence of necrosis, and distance to any putative intracerebral lesion). In addition, samples from tumor cases were neuropathologically reviewed to rule out the presence of tumor cells in the examined neocortical specimen.

### Slice preparation

Rats were deeply anaesthetized with isoflurane, decapitated, and the brain was quickly removed. Human cortical tissue was carefully micro-dissected and resected with minimal use of bipolar forceps to ensure tissue integrity. Rat and human neocortical tissue blocks were directly transferred into an ice-cold artificial cerebrospinal fluid (ACSF), which was either sucrose- or choline-based, respectively. The choline-based ACSF contained (in mM): 110 choline chloride, 26 NaHCO_3_, 10 D-glucose, 11.6 Na-ascorbate, 7 MgCl_2_, 3.1 Na-pyruvate, 2.5 KCl, 1.25 NaH_2_PO_4_,, and 0.5 CaCl_2_) (325 mOsm/l, pH 7.45).

To dissect human brain tissue, the pia was carefully removed using forceps and the pia-white matter (WM) axis was identified. 300 µm or 350 µm thick slices were prepared using a Leica VT1200 vibratome in ice-cold sucrose-based ACSF solution containing (in mM): 206 sucrose, 2.5 KCl, 1.25 NaH_2_PO_4_, 3 MgCl_2_, 1 CaCl_2_, 25 NaHCO_3_, 12 N-acetyl-L-cysteine, and 25 glucose (325 mOsm/l, pH 7.45). During slicing, the solution was constantly bubbled with carbogen gas (95% O_2_ and 5% CO_2_). After cutting, slices were incubated for 30 min at 31–33°C and then at room temperature in ACSF containing (in mM): 125 NaCl, 2.5 KCl, 1.25 NaH_2_PO_4_, 1 MgCl_2_, 2 CaCl_2_, 25 NaHCO_3_, 25 D-glucose, 3 myo-inositol, 2 sodium pyruvate, and 0.4 ascorbic acid (300 mOsm/l; 95% O_2_ and 5% CO_2_). To maintain adequate oxygenation and a physiological pH level, slices were kept in carbogenated ACSF (95% O_2_ and 5% CO_2_) during transportation.

### Whole-cell recordings

Whole cell recordings were performed in acute slices of rat and human neocortex. The recordings were conducted within 8 hours after slice preparation for rat neocortex and 30 hours at most for human neocortical tissue. During whole-cell patch-clamp recordings, rat or human slices were continuously perfused (perfusion speed “5 ml/min) with ACSF bubbled with carbogen gas and maintained at 30–33°C. Patch pipettes were pulled from thick wall borosilicate glass capillaries and filled with an internal solution containing (in mM): 135 K-gluconate, 4 KCl, 10 HEPES, 10 phosphocreatine, 4 Mg-ATP, and 0.3 GTP (pH 7.4 with KOH, 290–300 mOsm). Neurons were visualized using either Dodt gradient contrast or infrared differential interference contrast microscopy. In rat acute prefrontal cortical slices, cortical layers were distinguished by cell density and soma size, in accordance with previous studies on the PFC [40–43]. The PFC can be divided into three sections: the upper third comprises L1–L3, the middle third L5 and the lower third L6. Human L6 neurons were identified and patched according to their somatic location [44]. Putative excitatory neurons and interneurons were differentiated based on their intrinsic action potential (AP) firing pattern during recording and their morphological appearance after post hoc histological staining. Interneurons were excluded from the analysis.

Whole-cell patch clamp recordings of human or rat L6 neurons were made using an EPC10 amplifier (HEKA). During recording, slices were perfused in ACSF at 31–33°C. Signals were sampled at 10 kHz, filtered at 2.9 kHz using Patchmaster software (HEKA), and later analyzed offline using Igor Pro software (Wavemetrics). Recordings were performed using patch pipettes of resistance between 5 and 10 MΩ. Biocytin was added to the internal solution at a concentration of 3–5 mg/ml to stain patched neurons. A recording time >15 min was necessary for an adequate diffusion of biocytin into dendrites and axons of patched cells [45].

### Drug application

Acetylcholine was bath applied (10 µM) for 150–300 s through the perfusion system or puff applied (100 µM or 1 mM) during whole-cell patch clamp recordings. To specifically investigate the nicotinic effect of ACh, 200 nM atropine was bath applied to block the muscarinic responses. Tetrodotoxin (TTX, 0.5 µM) was added into perfusion ACSF for puff-application experiments to block AP firing induced by high concentrations of ACh. The puff pipette was placed at 10–20 µm from the recorded neuron, and a brief low pressure was applied for about 1 s. When different concentrations of ACh were administered to the same neuron, a second puff pipette from the same pulling pair was replaced to ensure uniformity in the electrode’s opening size. Galantamine (1 µM) was used as a PAM of α_5_*nAChRs. Drugs were purchased from Sigma-Aldrich or Tocris.

### Histological staining

After recordings, brain slices containing biocytin-filled neurons were fixed for at least 24 h at 4 °C in 100 mM phosphate buffer solution (PBS, pH 7.4) containing 4% paraformaldehyde (PFA). After rinsing several times in 100 mM PBS, slices were treated with 1% H_2_O_2_ in PBS for about 20 min to reduce any endogenous peroxidase activity. Subsequently, slices were rinsed repeatedly with PBS and then incubated in 1% avidin-biotinylated horseradish peroxidase (Vector ABC staining kit, Vector Lab. Inc.) containing 0.1% Triton X-100 for 1 h at room temperature. The reaction was catalyzed using 0.5 mg/ml 3,3-diaminobenzidine (DAB; Sigma-Aldrich) as a chromogen. Subsequently, slices were rinsed with 100 mM PBS, followed by slow dehydration with ethanol in increasing concentrations, and finally in xylene for 2–4 h [45]. After that, slices were embedded using Eukitt medium (Otto Kindler GmbH).

### Morphological 3D reconstructions and spine counting

Using NEUROLUCIDA® software (MBF Bioscience, Williston, VT, USA), morphological three-dimensional reconstructions of biocytin filled rat and human L6 excitatory neurons were made at a magnification of 1000-fold (100-fold oil-immersion objective and 10-fold eyepiece) on an upright microscope. Neurons were selected for reconstruction based on the quality of biocytin labelling when background staining was minimal. Neurons with major dendritic truncations due to slicing were excluded. Embedding with Eukitt medium reduced fading of cytoarchitectonic features and enhanced contrast between layers [45]. This allowed the reconstruction of different layer borders along with the neuronal reconstructions. Furthermore, the position of soma and layers were confirmed by superimposing the Dodt gradient contrast or differential interference contrast images taken during the recording. The tissue shrinkage was corrected using correction factors of 1.1 in the x–y direction and 2.1 in the z direction [45]. The spine counting was done using NEUROLUCIDA® software (MBF Bioscience, Williston, VT, USA). All dendritic spines were counted for at least 100 µm along a branch of both apical dendritic tuft and basal dendrite for each neuron. The spine density analysis was performed by using NEUROEXPLORER® software (MBF Bioscience, Williston, VT, USA).

### Data analysis

#### Whole-cell recording data analysis

Custom written macros for Igor Pro 6 (WaveMetrics) were used to analyze the recorded electrophysiological signals. The resting membrane potential (V_m_) of the neuron was measured directly after breakthrough to establish the whole-cell configuration with no current injection. The input resistance was calculated as the slope of the linear fit to the current–voltage relationship. For the analysis of single spike characteristics such as threshold, amplitude and half-width, a step size increment of 10 pA for current injection was applied to ensure that the AP was elicited very close to its rheobase current. The spike threshold was defined as the point of maximal acceleration of the membrane potential using the second derivative (d^2^V/dt^2^), which is, the time point with the fastest voltage change. The spike amplitude was calculated as the difference in voltage from AP threshold to the peak during depolarization. The spike half-width was determined as the time difference between rising phase and decaying phase of the spike at half-maximum amplitude.

The spontaneous activity was analyzed using the program SpAcAn (https://www.wavemetrics.com/project/SpAcAn). A threshold of 0.2 mV was set manually for detecting EPSP events.

#### Dendritic morphology and spine analysis

The horizontal dendritic fieldspan was defined as the widest distance between apical or basal dendrites, measured parallel to the pia. The vertical dendritic fieldspan was defined as the longest distance between apical tuft and basal dendrite, measured perpendicular to the pia. The aspect ratio of the dendritic fieldspan was calculated by dividing the vertical fieldspan by the horizontal fieldspan. Other morphological parameters such as total dendritic length, number of basal dendrites, soma area and spine density were analyzed with NEUROEXPLORER® software (MBF Bioscience, Williston, VT, USA).

#### Statistical analysis

Data was either presented as box plots (n ≥ 10) or as bar histograms (n < 10). For box plots, the interquartile range (IQR) is shown as box, the range of values within 1.5#IQR is shown as whiskers and the median is represented by a horizontal line in the box; for bar histograms, the mean % SD is given. Wilcoxon Mann-Whitney U test was performed to assess the difference between individual groups. To assess the differences between two paired groups under different pharmacological conditions, Wilcoxon signed-rank test was performed. Statistical significance was set at P < 0.05, and n indicates the number of neurons/slices analyzed.

## Results

### Comparing Electrophysiology and Morphology of mPFC L6 RS Neurons in WT, α5SNP and α5KO Rats

To investigate whether the expression of the α5 nAChR subunit affects the electrophysiological characteristics of L6 neurons in medial prefrontal cortex (mPFC), we prepared acute brain slices from rats that were either wild-type (WT), carrying the human coding polymorphism rs16969968 in *Chrna5* (α5SNP), or constitutively lacking the α5 subunit (α5KO) [39] (**Figure 1A**). L6 excitatory neurons comprise various electro-morphological subtypes, and neurons showing different firing patterns also exhibit distinct passive properties [46–49]. Here we focused on analyzing the intrinsic properties of the L6 regular spiking (RS) excitatory neurons (**Figure 1C**), which are considered to be corticothalamic (CT) projecting neurons and are known to express the α5 nAChR subunit encoded by *Chrna5* [21, 50, 51]. L6 RS neurons in α5KO rats exhibited the most depolarized resting membrane potential (Vrest, -65.0 % 7.0 mV) among the three rat genotypes (-71.7 % 4.2 mV for WT and -69.7 % 4.0 mV for α5SNP). Correspondingly, L6 RS neurons in α5KO rats demonstrated significantly increased excitability, as evidenced by smaller rheobase currents compared to the WT (58.4 % 29.3 vs. 87.7 % 24.9 pA, **P < 0.01) and α5SNP group (58.4 % 29.3 pA vs. 86.0 % 29.1 pA, *P < 0.05), respectively (**Figure 1B, F**). The input resistance (R_in_) of L6 RS neurons in the α5KO group was significantly higher than R_in_ of neurons in WT (330 ± 60 vs. 251 ± 44, ***P < 0.001) or α5SNP (330 ± 60 vs. 255 ± 70, **P < 0.01) rats (**Figure 1D, F**). In the three genotypes, we observed differences in the voltage sag following a hyperpolarizing current injection. L6 RS neuros in the α5KO groups exhibited the most prominent voltage sag (V_sag_, 1.65 ± 1.82 mV), which was more than twice that of neurons in the WT (0.70 ± 0.69 mV) or α5SNP (0.65 ± 0.34 mV) group (**Figure 1E, F**). Furthermore, neurons in the α5KO group exhibited a significantly smaller action potential (AP) amplitude compared to those in the WT group (89.8 ± 6.4 vs. 94.5 ± 5.2 mV, *P < 0.05). Additionally, they displayed a significantly longer onset time (339 ± 129 vs. 219 ± 173 ms, *P < 0.05) for the first AP evoked by injecting a rheobase current. In contrast to the increased excitability observed in L6 RS neurons of the α5KO group, no significant differences were observed in the electrophysiological properties between neurons in the WT and α5SNP groups (**Table S1**). This suggests that the deletion of *Chrna5* has an impact on the electrophysiological characteristics of L6 RS neurons in the rat mPFC, while those in rats expressing the human SNP remain unaffected.

**Figure 1.**
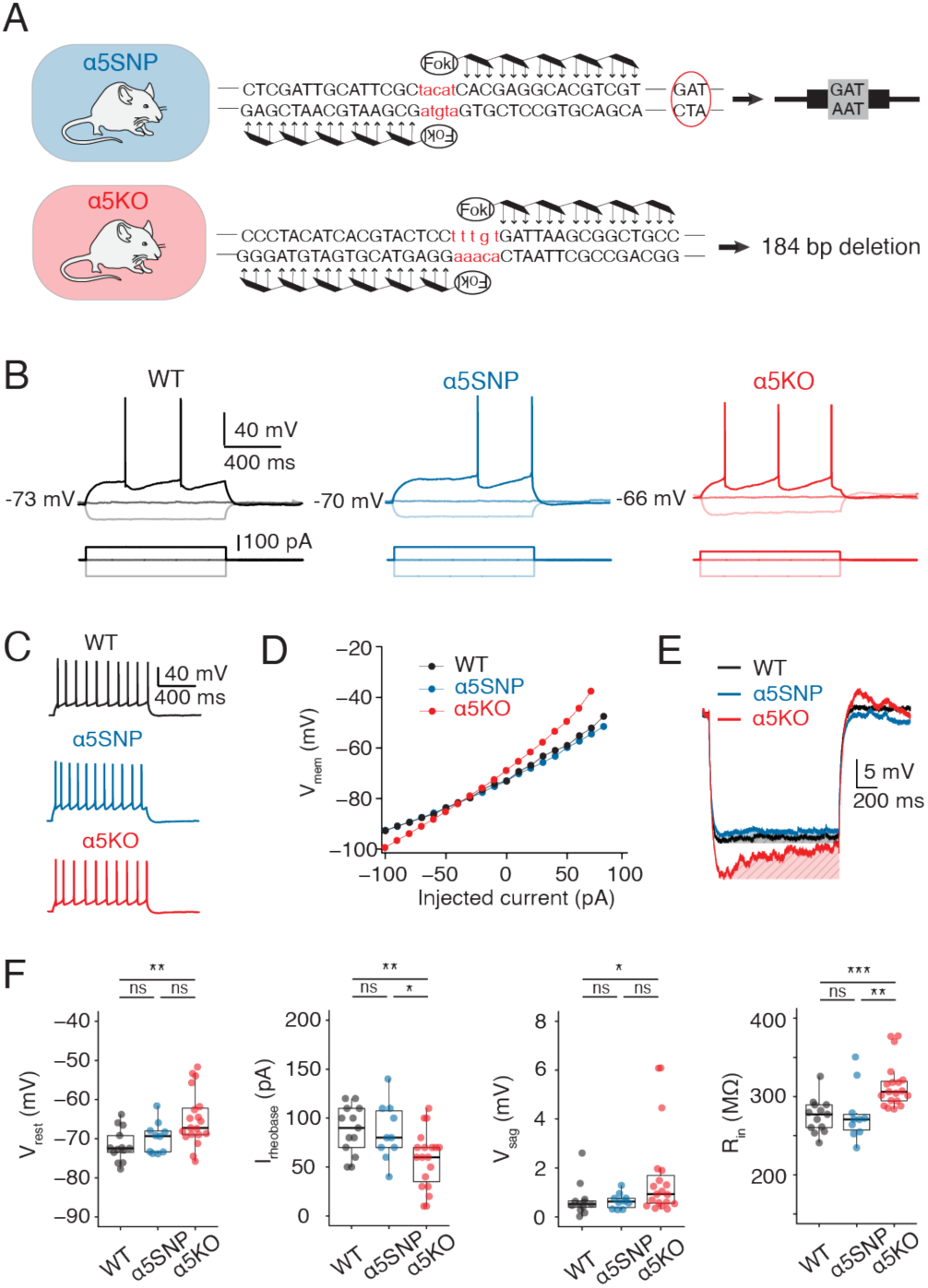
L6 regular spiking (RS) neurons in *α*5KO rats show higher intrinsic excitability than those in WT and *α*5SNP rats. **(A)** Top, creation of α5SNP rats showing the insertion of the α5SNP at the end of the exon 5 of *Chrna5*. Bottom, creation of α5KO rats showing a 184-bp deletion in the gene *Chrna5*. **(B)** Representative AP firing of a L6 RS neuron in a WT, α5SNP and α5KO rat. AP firing (top) was elicited by a 1 s square current of −100 pA, 0 pA and rheobase current (bottom). **(C)** Representative firing patterns of the same L6 RS neurons shown in B. **(D)** I–V relationship of the same L6 RS neurons shown in B showing that the neuron in α5KO rat display a larger membrane input resistance than the other genotypes. **(E)** Voltage response to a hyperpolarizing step current of −100 pA and duration of 1 s in the same L6 RS neurons shown in B. The red shaded area shows the amplitude of voltage sag. **(F)** Summary data of several electrophysiological properties of L6 RS neurons in WT, α5SNP and α5KO rats. Data were compared between WT (n = 13), α5SNP (n = 10) and α5KO (n = 19) groups and were presented as box plots,*P < 0.05, **P < 0.01,***P < 0.001 for the Wilcoxon Mann– Whitney U test; ns, not significant.

It has been shown that the α5 subunit plays an important role in normal developmental changes of the dendritic morphology of L6 pyramidal cells. These differences in dendritic morphology can be observed from early adulthood between WT and α5KO mice [52]. To study the impact of the rs16969968 polymorphism on neuronal morphology, we reconstructed the 3D somato-dendritic morphology of L6 RS neurons in the mPFC of young adult WT, α5SNP and α5KO rats. As shown in **Figure 2A**, apical dendrites of L6 RS neurons in WT group often terminate before reaching cortical layer 1 (L1, 55%). In contrast, all α5SNP and 80% α5KO rats exhibit long apical dendrites extending into L1 of the mPFC (**Figure 2A, B**). There is a gradual decrease of the horizontal dendritic fieldspan that correlates with the reduced presence of functional α5*nAChRs and therefore a gradual increase in the aspect ratio of the dendritic fieldspan across the WT, α5SNP to α5KO groups (**Figure 2A, C**). Neurons in both α5SNP and α5KO groups showed a larger number of basal dendrites compared to those in WT rats. However, due to a markedly narrower horizontally dendritic arborization of neurons in the α5KO group, L6 α5SNP neurons exhibited a greater total length of basal dendrites compared to neurons in the other two genotypes (**Figure 2C**, **Table S1**). More details regarding morphological properties and statistical comparisons are given in **Supplementary Table S1**. Our findings suggest that morphological changes in L6 RS neuron dendrites, which depend on the α5 subunit, are evident in rats carrying the SNP.

**Figure 2.**
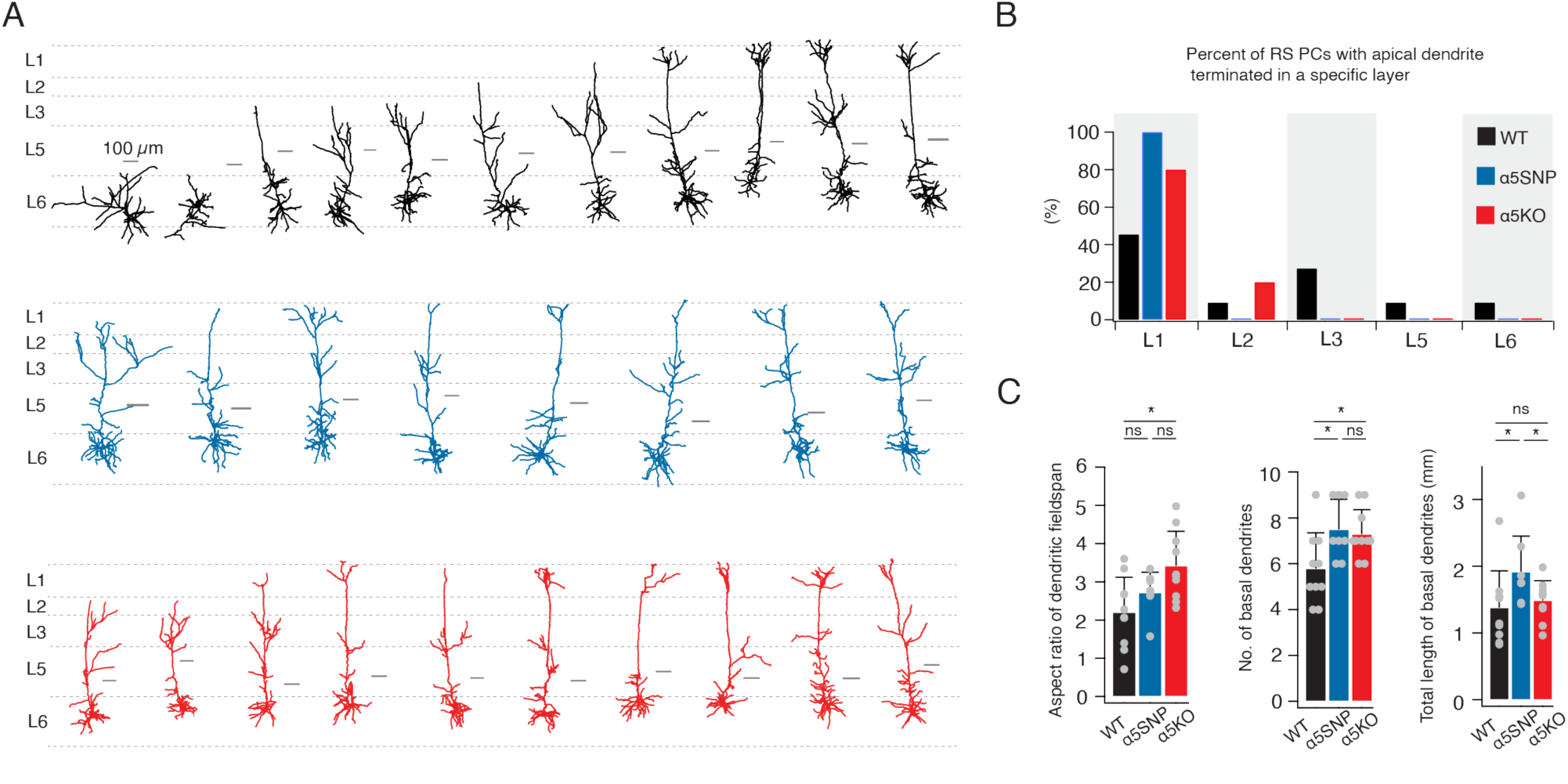
Comparison of dendritic morphology of L6 RS neurons between WT, *α*5SNP and KO genotypes. **(A)** Individual dendritic reconstruction of L6 RS neurons in the three rat genotypes. Neurons are shown in their laminar location with respect to averaged cortical layers, scale bar of each individual reconstruction is given. L6 RS neurons of WT (n = 11) rats are shown in black, α5SNP (n = 8) in blue and α5KO (n = 10) in red. **(B)** Histograms of the percentage of L6 RS neurons with apical dendrite terminated in each cortical layer. L6 RS neurons of WT (n = 11) rats are shown in black, α5SNP (n = 8) in blue and α5KO (n = 10) in red. **(C)** Summary data of several morphological properties of L6 RS neurons in WT, α5SNP and α5KO rats. Data were compared between WT (n = 11), α5SNP (n = 8) and α5KO (n = 10) groups, *P < 0.05 for the Wilcoxon Mann–Whitney U test; ns, not significant.

### Functional Presence of α5 nAChR Subunit Correlates with Dendritic Spine Density of L6 RS Neurons

Previous studies revealed that dysfunction of high affinity nicotinic receptors or long term exposure of nicotinic agonist have an impact on dendritic spine density [53–56]. This phenomenon is also observed in L6 pyramidal cells in the PFC of mice lacking the α5 subunit in nicotinic receptors [37]. Here we analyzed the spine density in distal apical tuft as well as basal dendrite near the soma of L6 RS neurons in the three rat genotypes. Our results revealed that the complete deletion of the α5 nAChR subunit led to a substantial reduction in the spine density of apical tuft dendrites (0.21 ± 0.04 vs. 0.41 ± 0.10 µm^-1^, ***P < 0.001). In contrast, neurons in the SNP group exhibited only a modest and statistically not significant decrease in the spine density of the apical dendrite (0.36 ± 0.09 vs. 0.41 ± 0.10 µm^-1^, P = 0.514) (**Figure 3A**). Furthermore, dysfunction of the α5 subunit also led to a gradual reduction in the basal dendritic spine density. In comparison to L6 excitatory neurons in WT rats, those in α5SNP and α5KO rats exhibited a significant decrease in basal dendritic spine density, namely 0.38 ± 0.09 µm^-1^ (*P < 0.05) and 0.30 ± 0.10 µm^-1^ (***P < 0.001), respectively, compared to 0.48 ± 0.07 µm^-1^ in WT rats (**Figure 3B**).

**Figure 3.**
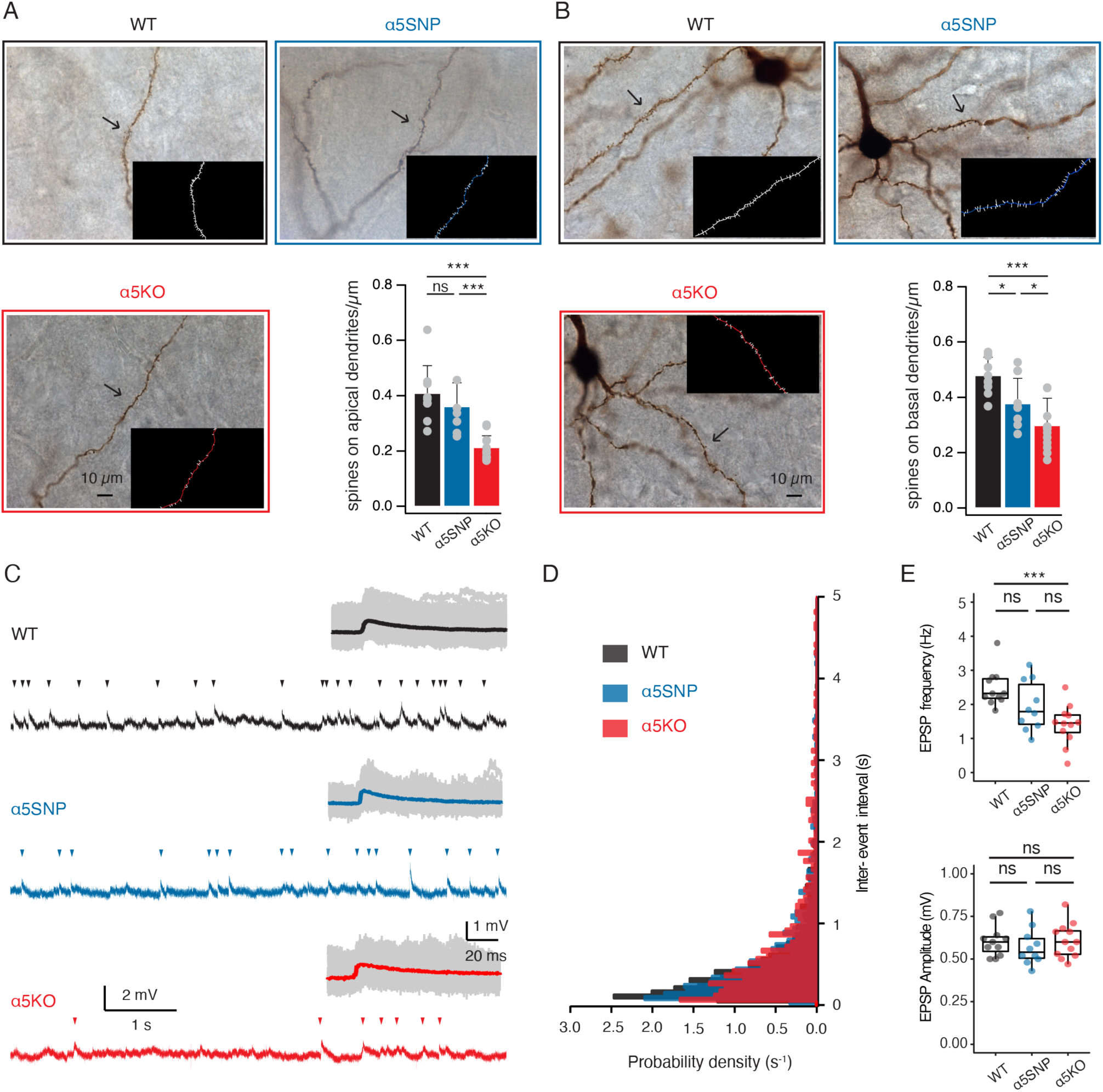
Down-regulation or depletion of *α*5 subunit confers a decrease of dendritic spine density and EPSP frequency in L6 RS neurons. **(A)** Photomicrographs of biocytin-filled apical tuft of a representative L6 RS neuron in a WT, α5SNP and α5KO rat. Insets showing counted dendritic spines of marked branch. Spine density on apical tuft was compared between neurons from WT (n = 8), α5SNP (n = 7) and α5KO (n = 10) groups, ***P < 0.001 for the Wilcoxon Mann–Whitney U test; ns, not significant. **(B)** Photomicrographs of biocytin-filled basal dendrites of a representative L6 RS neuron in a WT, α5SNP and α5KO rat. Insets showing counted dendritic spines of marked branch. Spine density on basal dendrite was compared between neurons from WT (n = 8), α5SNP (n = 7) and α5KO (n = 10) groups, *P < 0.05, ***P < 0.001 for the Wilcoxon Mann–Whitney U test. **(C)** A 7 s current-clamp recording of a L6 RS neuron in a WT, α5SNP and α5KO rat. Excitatory postsynaptic potentials (EPSPs) are marked by arrowheads. Insets displaying the overlay of EPSPs extracted from a 20 s continuous recording of the same neurons. The average and individual EPSPs are superimposed and given in a color and gray shade, respectively. **(D)** Histograms showing probability density of inter-event intervals between two consecutive EPSPs. Data were collected and analyzed from 50s continuous recordings of 11 L6 RS neurons in WT rats (n = 1299 events), 10 neurons in α5SNP rats (n = 828 events) and 12 neurons in α5KO rats (n = 622 events). **(E)** Box plots comparing EPSP frequency and amplitude of L6 RS neurons in WT (n = 11 neurons), α5SNP (n = 10 neurons) and α5KO rats (n = 12 neurons); *** P < 0.001 for the Wilcoxon Mann–Whitney U test; ns, not significant.

To investigate whether the decrease in spine density resulted in a reduced spontaneous synaptic activity, we determined the frequency and amplitude of spontaneous excitatory postsynaptic potentials (sEPSPs) during continuous, 50 s long current-clamp recordings from L6 RS neurons (**Figure 3C, D**). In parallel with a reduced functional expression of the α5 subunit, there was a gradual decrease in EPSP frequency. Neurons in the α5KO group showed a much lower EPSP frequency compared to WT neurons (1.40 ± 0.58 vs. 2.47 ± 0.54 Hz, ***P < 0.001) while α5SNP neurons displayed an intermediate value between the other two genotypes (**Figure 3E**). Additionally, there was no discernible difference in the amplitude, decay time or rise time of spontaneous EPSPs among neurons in the WT, α5SNP, and α5KO groups (**Figure 3C, E**). The data indicates that L6 RS neurons in mPFC of three rat genotypes differ significantly in spine density on their apical and basal dendrites, which is reflected in their spontaneous synaptic activity.

### Galantamine Acts as a Positive Allosteric Modulator of α5*nAChRs in L6 RS Neurons

To examine the nAChR responses in L6 of rat mPFC in isolation, we blocked muscarinic AChRs (mAChRs) by bath-applying 200 nM atropine in the perfusion ACSF and performed whole-cell recordings from L6 RS neurons. Following application of 10 µM acetylcholine (ACh) for 60 seconds, L6 RS neurons in WT rats showed an average membrane potential depolarization of 3.1 ± 3.2 mV. The vast majority of L6 RS neurons in α5SNP and α5KO rats showed weak to no depolarization following ACh application, as suggested by an average membrane potential change of 0.77 ± 0.73 mV and 0.93 ± 1.04 mV, respectively (**Figure 4A-C**). Galantamine is a positive allosteric modulator (PAM) of α5*nAChRs at lower concentrations and also an inhibitor of acetylcholinesterase [23, 50]. To test whether galantamine exerts an allosteric modulation of nicotinic responses in L6 neurons, we pre-applied 1 µM galantamine for 10 minutes before and concurrently with ACh application. In the presence of galantamine, we observed a ∼3.5-fold enhancement of the nicotinic ACh responses in the WT group, *i.e*. from 3.1 % 3.2 to 11.7 % 6.2 mV (***P < 0.001). In α5SNP rat neurons, galantamine significantly increased ACh-induced depolarization from 0.8 % 0.7 to 5.5 % 4.7 mV (***P < 0.001), leading to nicotinic responses comparable to the control level observed in WT neurons without the galantamine application (5.5 % 4.7 vs. 3.1 % 3.2 mV, P = 0.11) (**Figure 4B,C**). This implies that galantamine functions as a PAM of α5*nAChRs and can restore the dysfunctional nAChR responses in L6 neurons of α5SNP rats to normal levels. In α5KO rat neurons, we found no statistically significant difference in ACh-induced responses before and after galantamine treatment (0.9 % 1.0 vs. 1.2 % 1.7 mV, P = 0.679), suggesting a specific modulation of galantamine on α5*nAChRs (**Figure 4B,C**).

**Figure 4.**
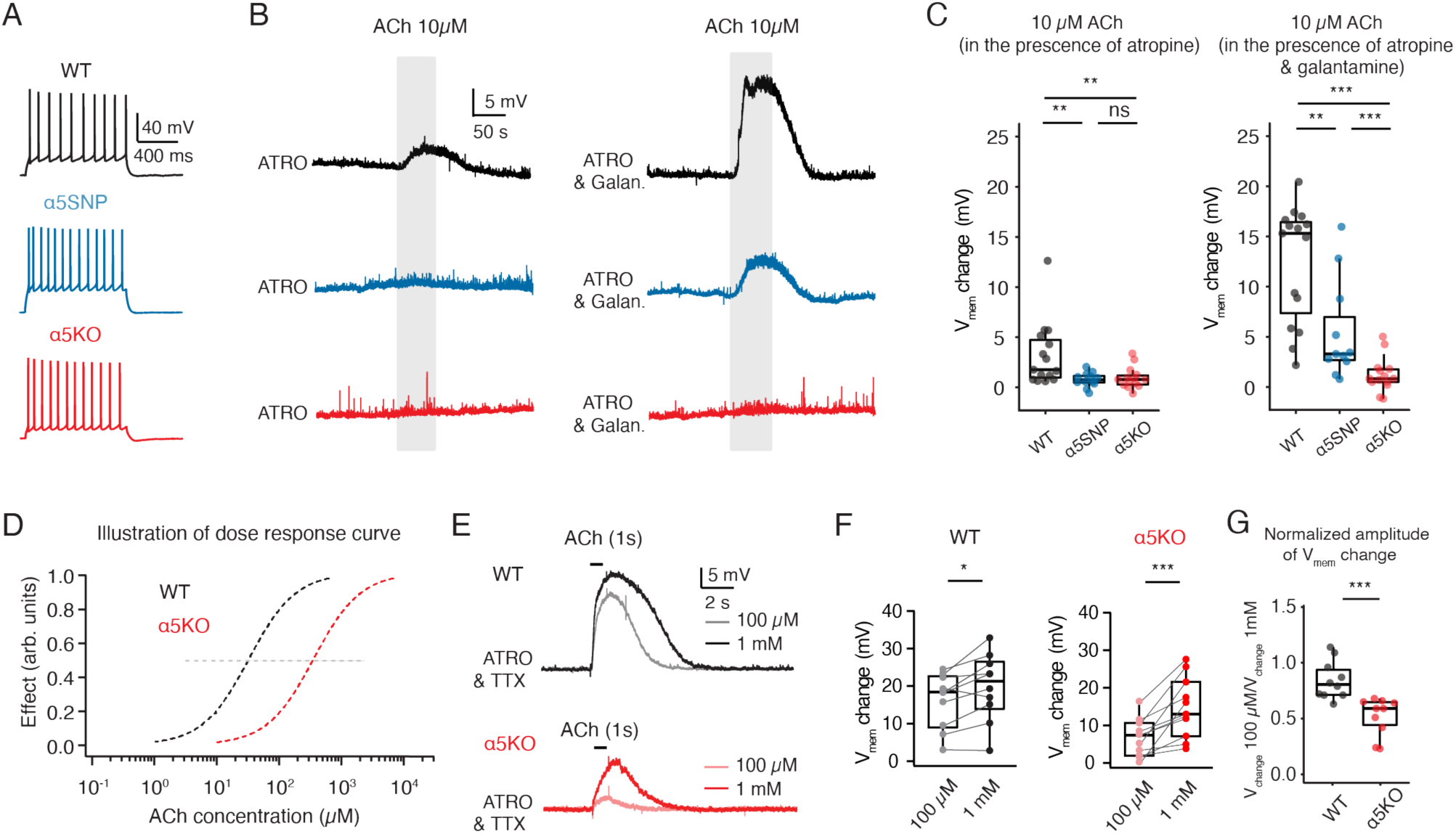
Galantamine acts as a positive allosteric modulator of *α*5*nAChRs in L6 RS neurons. **(A)** Representative firing patterns of a L6 RS neuron recorded in WT α5SNP and α5KO rat, respectively. **(B)** Bath application of a low concentration of ACh (10 µM, 50 s) to L6 RS neurons was enhanced in the presence of 1 µM galantamine in WT and α5SNP rats, but not in α5KO rats, compared to the absence of galantamine. Recordings were performed in the prescence of 200 nM atropine to rule out the muscarinic receptor activation. Representaive recording traces are shown in balck for WT neuron, blue for α5SNP neuron and red for α5KO neuron, respectively. **(C)** Summary box plots showing the ACh (10 µM) induced resting membrane potential (Vm) change under control and galantamine (1 µM) conditions in L6 RS neurons in WT (n = 16 neurons), α5SNP (n = 12 neurons) and α5KO rats (n = 15 neurons). **P < 0.01, ***P < 0.001 for the Wilcoxon Mann–Whitney U test; ns, not significant. **(D)** As a consequnce of deletion of α5 subunits,ACh activate nAChRs of L6 RS neurons in WT rats at significantly lower concentrations than in α5KO rats, as can be seen from the illustration of dose–response curves; the grey dashed line marks the EC50 for the curves. **(E)** Representative traces of puff-applied ACh at 100 µM and 1 mM concentrations for 1 s in L6 RS neuron in a WT (top, black) and α5KO (bottom, red) rat. Recordings were performed in the prescence of 200 nM atropine and 0.5 µM tetrodotoxin (TTX) to block mAChRs activation and AP firing. ACh-evoked Vm depolarization is smaller in KO rat but shows a substantial increase when using high concentration of 1 mM ACh when compared to 100 µM. **(F)** Box plots of ACh-evoked (1s puff-applied) depolarizations of L6 RS neurons under 100 µM and 1 mM concentrations of ACh in WT (n = 10 neurons) and α5KO (n = 11 neurons) rats. *P < 0.05, ***P < 0.001 for Wilcoxon signed-rank test. **(G)** Normalized amplitude of Vm change (Vchange 100µM/Vchange 1mM) showing that a significant difference in dose-dependent responses between L6 RS neurons in WT (n = 10 neurons) and α5KO (n = 11 neurons) rats.

Previous studies have demonstrated that α4β2*nAChRs locate specifically within mPFC L6 neurons [50, 57, 58], which is one of the known type of nAChRs that potentially includes α5 subunits as part of their assemblies [59]. The co-assembly of α5 subunits into α4β2*nAChRs enhances its Ca^2+^ permeability and slows down the receptor desensitization [22]. It has been shown that the deletion of the α5 subunit does not affect the density of α4β2 nAChRs but specifically reduces agonist activation affinity [52, 60]. Therefore we hypothesized that as a consequence of α5 subunit deletion, L6 neurons in the α5KO group require higher concentrations of ACh for activating nAChRs (**Figure 4D**). To test this hypothesis, we puff-applied high concentrations of ACh (100 µM and 1 mM) for 1 second sequentially on L6 RS neurons in WT and α5KO rats. Recordings were performed in the presence of 200 nM atropine and 0.5 µM tetrodotoxin (TTX) to block mAChRs and AP firing. Neurons in WT rats showed consistently strong depolarizations with a slight increase from 16.2 % 7.3 to 19.8 % 8.9 mV (*P < 0.05), in response to the puff-application of 100 µM and 1 mM ACh, respectively. In contrast, neurons in α5KO rats displayed a much greater depolarization following the application of 1 mM ACh compared to 100 µM ACh (14.7 % 7.9 vs. 6.8 % 4.9 mV ***P < 0.001) (**Figure 4E,F**). A significant difference in the normalized amplitude of membrane potential change induced by 100 µM ACh (normalized to the effect of 1 mM ACh) was observed between L6 WT and α5KO neurons (**Figure 4G**). In summary, dysfunction of the α5 nAChR subunit resulted in a shift of the ACh dose-response curve to higher concentrations, suggesting a potential mechanism underlying nicotine dependence.

### Selective Functional Expression of the α5 nAChR Subunit in L6 RS rather than BS Neurons

In mPFC L6, a distinct population of neurons displaying burst-spiking (BS) firing patterns was identified in addition to regular spiking neurons. While L6 corticothalamic neurons display a regular AP firing pattern, L6 corticocortical (CC) neurons tend to exhibit burst-spiking behavior [46, 47, 49, 61]. These neurons can be easily distinguished from RS neurons by their burst-like spiking pattern consisting of two or three initial, closely spaced spikes (**Figure 5A**). This is evident by a smaller second AP amplitude and a reduced adaptation ratio of the second versus tenth inter-spike intervals (ISI2/ISI10) compared to those of RS neurons (**Figure 5A,B**). Notably, BS neurons display significantly weaker nicotinic responses compared to RS neurons in L6 of both rat barrel cortex and human mPFC [19, 50]. To examine whether this is due to the absence of α5 subunit in BS neurons, we puff-applied 100 µM ACh for 1 second on L6 neurons in WT and α5KO rats. Recordings were conducted in the presence of 200 nM atropine and 0.5 µM tetrodotoxin (TTX) to exclude the influence of mAChR activation and AP firing. Consistent with previous findings, BS neurons showed a much smaller ACh-induced nicotinic response compared to RS neurons (4.4 % 4.3 vs. 16.0 ± 7.5 mV, *P < 0.05) in WT rats (**Figure 5C,D**). In α5KO rats, the deletion of the α5 nAChR subunit resulted in a decreased nicotinic response in RS neurons (6.1 ± 5.2 vs. 16.0 ± 7.5 mV, *P < 0.05). However, the amplitude of the ACh-induced depolarization in BS neurons remained unaffected compared to that in WT rats (2.3 ± 3.1 vs. 4.4 ± 4.3 mV, P = 0.352). This suggests that, there is an absence of α5 subunit co-assembly in nAChRs on BS neurons, unlike RS neurons. It is worth noting that even in α5KO rats, BS neurons showed a smaller nicotinic response compared to RS neurons (2.3 ± 3.1 vs. 6.1 ± 5.2 mV, *P < 0.05) (**Figure 5D,E**). This suggests a lower density of functional nAChR presence in L6 BS neurons, at least at the soma.

**Figure 5.**
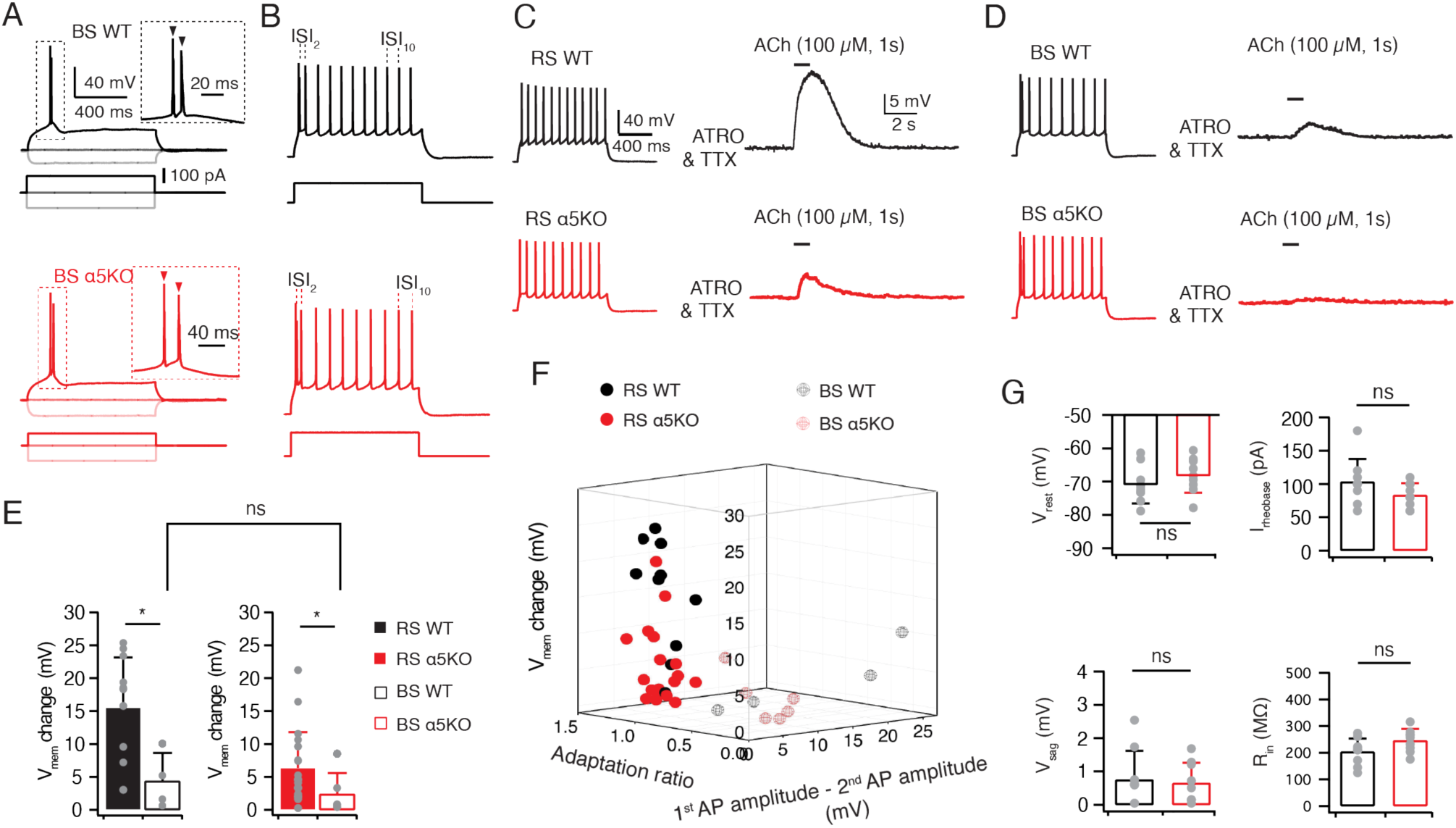
*α*5 subunits are abundantly expressed in L6 RS neurons but not in busrt spiking (BS) neurons. **(A)** Representative AP firing of a L6 BS neuron in a WT, α5SNP and α5KO rat. AP-firing (top) was elicited in response to 1 s square current of −100 pA, 0 pA and rheobase current (bottom). Insets showing initial AP bursts at an expanded time scale. **(B)** Corresponding firing patterns of representative BS neurons shown in A. Injected current pulses are shown at the bottom. **(C)** Representative traces of puff-applied 100 µM ACh for 1 s in a L6 RS neuron in a WT (top, black) and α5KO (bottom, red) rat. Recordings were performed in the prescence of 200 nM atropine and 0.5 µM tetrodotoxin (TTX) to block mAChR activation and AP firing. ACh-evoked depolarizations are smaller in α5KO compared to WT rat. The corresponding firing patterns of the neurons are shown on the left. **(D)** Representative traces of puff-applied 100 µM ACh for 1 s in a L6 BS neuron in a WT (top, black) and α5KO (bottom, red) rat. Recordings were performed in the prescence of 200 nM atropine and 0.5 µM tetrodotoxin (TTX) to block mAChR activation and AP firing. The ACh-evoked depolarization is small both in WT and α5KO rat. The corresponding firing patterns are shown on the left. **(E)** Summary histograms showing that puff application of ACh (100 µM) induces smaller depolarization in L6 BS neurons than RS neurons in both WT (n = 14 for RS neurons and n = 4 for BS neurons) and α5KO rats (n = 23 for RS neurons and n = 5 for BS neurons). *P < 0.05 for the Wilcoxon Mann–Whitney U test; ns, not significant. **(F)** 3D scatter plot indicate a clear separation of L6 RS and BC neurons subtypes based on their electrophysiological properties and ACh-induced Vm changes. Neurons recorded from WT rats are shown in black and those from α5KO rats in red. **(G)** Histograms of several electrophysiological properties of L6 BS neurons between WT (n = 9 neurons, black) and α5KO (n = 9 neurons, red) rats. Statistical analysis was performed using the Wilcoxon Mann–Whitney U test; ns, not significant.

We analyzed the electrophysiological properties of BS neurons in WT and α5KO rats. In **Figure 5 F**, ACh-induced membrane potential changes were plotted against the difference between the first and second AP amplitude, as well as the AP adaptation ratio. The 3D scatter plot revealed a strong correlation between cholinergic response and firing pattern-related properties for the two distinct types of excitatory neurons in L6. Moreover, the deletion of the α5 subunit in α5KO rats did not affect the electrophysiological properties of BS neurons, including resting membrane potential, rheobase current, voltage sag, and input resistance, which were not significantly different between WT and α5KO BS neurons (**Figure 5G**). This lends further support to the idea that L6 BS excitatory neurons do not express the α5 nAChR subunit and hence only possess α4β2*nAChR.

### Cell Type-Specific Modulation of nAChRs by Galantamine in Human Neocortical Layer 6

For experiments on human L6 neurons, acute brain slices were prepared from tissue blocks obtained from brain surgery in six patients aged 23-66 years. The neocortical tissue used was from either the frontal, temporal or parietal cortex (**Supplementary Table S2**). It was resected during surgical access to the pathological brain region and was sufficiently distant from the pathological focus. Therefore, it can be considered as healthy cortex (**Figure 6A**). Whole-cell current clamp recordings with simultaneous biocytin filling were made from human cortical L6 neurons allowing post-hoc identification of their morphology. To analyze the repetitive AP firing properties of L6 neurons, voltage recordings were used in which a current injection elicited ∼10 APs. Similar to observations in rodents, two distinct firing patterns were identified in human L6 excitatory neurons: regular or burst spiking (**Figure 6B**). In contrast to human L6 RS neurons, which exhibited nearly constant AP amplitudes throughout the train, BS neurons often displayed a spike burst at the onset of the AP train, with a smaller second AP amplitude and subsequent recovery in AP magnitude (**Figure 6C**). Moreover, RS neurons exhibited either a single AP or a spike doublet that occurred at the beginning of the AP train and was followed by APs with nearly constant ISIs. In contrast, BS neurons displayed an initial long ISI but a shorter stable ISIs thereafter following the initial AP burst (**Figure 6C**). Notably, at the rheobase current injection, a spike doublet was only observed for BS but not RS neurons (**Figure 6B**). Significant differences in the adaptation ratio, voltage sag and frequency-current slope were found between human RS and BS excitatory neurons (**Figure 6C** and **Supplementary Table S3**). More electrophysiological properties and the statistical comparison of the two neuron types are given in **Supplementary Table S3**.

**Figure 6.**
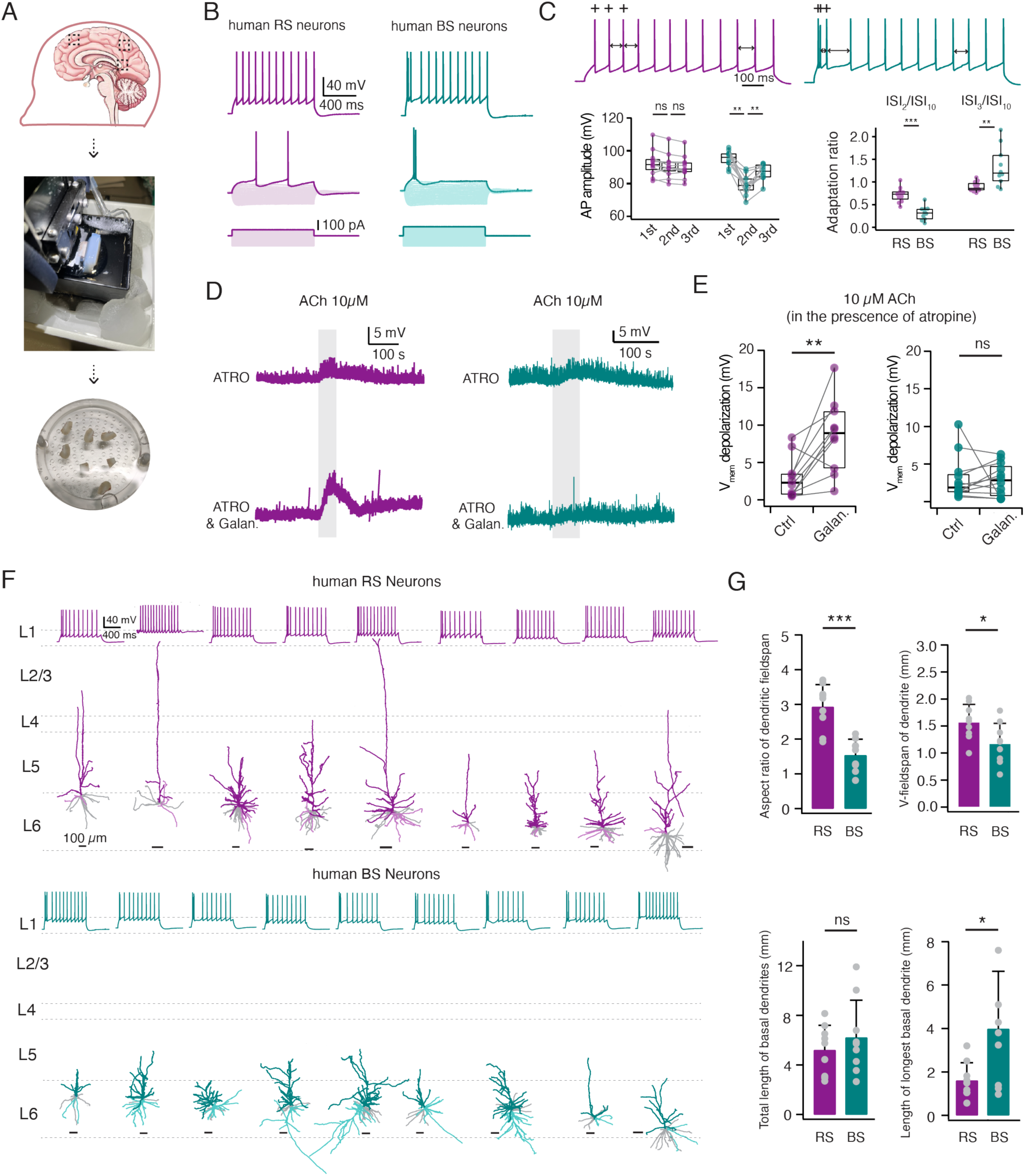
Glantamine acts as a positive allosteric modulator of nAChRs in human L6 RS but not BS neurons. **(A)** Summary procedure of acute human brain slice preparation. **(B)** Top, representative firing patterns of of a human L6 RS (left) and BS neuron (right). Bottom, AP-firing was elicited in response to 1 s square current using a step size increment of 10 pA from -100 pA to its rheobase current. **(C)** Top, representative firing patterns at an expanded time scale of the same human L6 RS and BS neurons shown in B. The first three APs are marked by +. The 2^nd^, 3^rd^, and 10^th^ inter-spike interval are marked by bidirectional arrow. Bottom left, box plots comparing the first, second and third AP amplitude of human L6 RS (n = 11) and BS (n = 11) neurons.**P < 0.01 for Wilcoxon signed-rank test; ns, not significant. Bottom right, box plots comparing spiking adaptation bwtween RS and BS neurons. **P < 0.01, ***P < 0.001 for Wilcoxon Mann–Whitney U test. **(D)** In the presence of 1 µM galantamine, bath application of a low concentration ACh (10 µM, 50 s) induces a larger depolarization of human L6 RS but not of BS neurons compared to the abscence of galantamine. Recordings were performed in the prescence of 200 nM atropine to block mAChR activation. Representative recordings are shown in purple for RS neurons while in teal for BS neurons. **(E)** Box plots showing that galantamine enhances ACh-induced Vm depolarization of human L6 RS (n = 10) neurons but affect not that of BS (n = 11) neurons. **P < 0.01 for Wilcoxon signed-rank test; ns, not significant. **(F)** Individual dendritic reconstruction of human L6 RS and BS neurons. Neurons are shown in their laminar location with respect to averaged cortical layers, scale bar of each individual reconstruction is given. Apical dendrites of RS neurons (n = 9) are shown in purple and BS neurons (n = 9) in teal. The longest basal dendrite of neurons are given in lighter shade and the other basal dendrites in gray. The top traces shows the corresponding firing pattern of each neuron. **(G)** Histograms comparing several dendritic properties between L6 RS (n = 9) and BS (n = 9) neurons. *P < 0.05, **P < 0.01 for Wilcoxon signed-rank test; ns, not significant.

To examine the nicotinic responses in L6 of human neocortex, we blocked mAChRs by bath-applying 200 nM atropine in the perfusion ACSF. Following bath-application of 10 µM ACh, both human L6 RS and BS neurons showed a membrane potential depolarization of 3.0 ± 2.8 mV and 3.1 ± 3.1 mV, respectively. To identify whether the ACh response can be potentiated, 1 µM galantamine was pre-applied for 10 minutes before the ACh application. The majority of the RS neurons showed a significant increase in the nicotinic response from 3.0 ± 2.8 to 8.6 ± 4.9 mV (**P < 0.01) following galantamine treatment, indicating the presence of α5*nAChRs. In contrast, the ACh response in L6 BS neurons was not changed by galantamine (3.1 ± 3.1 vs. 2.9 ± 2.3 mV, P = 0.884) (**Figure 6D,E**).

To compare the morphological differences between two types of L6 neurons, we performed 3D reconstructions of the somato-dendritic domain of RS and BS neurons in the human neocortex. RS neurons exclusively have an upright projecting apical dendrite that terminates within the range of cortical L1 to L5 while, BS neurons display various morphologies including small pyramidal cells, multipolar, or bipolar neurons (**Figure 6F**). On average, RS neurons showed a larger vertical dendritic fieldspan (1.6 ± 0.3 vs. 1.2 ± 0.4 mm, *P < 0.05), resulting in a greater aspect ratio of dendritic fieldspan when compared to BS neurons (2.9 ± 0.6 vs. 1.5 ± 0.5, ***P < 0.001).

Furthermore, a significant proportion of BS neurons exhibited a multipolar morphology (5 out of 9), so that on average, BS neurons showed a greater length of the longest basal dendrite compared to RS neurons (4.0 ± 2.6 vs. 1.6 ± 0.8 mm, *P < 0.05) (**Figure 6G**). Additional morphological properties and a statistical comparison of the two neuron types are presented in **Supplementary Table S3**.

## Discussion

In this study, we conducted whole-cell patch clamp recordings from L6 RS neurons in the mPFC of WT, α5SNP, and α5KO rats. We found that the deletion of the α5 subunit significantly altered the intrinsic membrane properties, spine density and dendritic morphology of L6 RS neurons. On the other hand, the presence of an α5SNP mutation affected spine distribution and dendritic arborizations without altering intrinsic electrophysiological properties. Furthermore, our findings revealed the crucial role of the α5 subunit encoded by *Chrna5* in generating nicotinic responses in response to low concentrations of ACh. Galantamine, a PAM of α5*nAChRs, effectively restored nicotinic modulation in neurons of α5SNP rats but not in α5KO rats. Additionally, we found that the functional distribution of the α5 subunit is cell-type specific, with a notably more prominent presence in RS neurons compared to BS neurons. This observation was consistent in both rat and human neocortical L6.

### Aberrant neuronal properties due to the loss or partial loss of function of the α5 subunit

Previous studies showed no significant differences in passive properties, including input resistance and resting membrane potential, between neurons in WT and α5KO mice. This observation applies to neurons located in both the mouse rostral interpeduncular nucleus and RS neurons in L6 of the mouse mPFC, which are two sites characterized by remarkably high *Chrna5* mRNA expression levels in rodents [58, 62]. In marked contrast, we observed that α5 deletion in the created KO rat line caused significant alterations in several electrophysiological characteristics of L6 RS neurons, including the resting membrane potential, input resistance, rheobase current and voltage sag. This may be due to species differences between transgenic mice and rats. We then analyzed the same properties of L6 RS neurons in mPFC of rats carrying the human α5SNP rs16969968. This genetic variant is highly prevalent in the general population, with a frequency of 37% in Europeans and 50% in Middle Eastern populations. It has been proven to be a risk factor for nicotine dependence and SZ [63–65]. In transgenic rodents, this α5SNP contributed to increased nicotine intake and the generation of Chronic Obstructive Pulmonary Disease (COPD)-like lesions [39, 66, 67]. *In vitro* data suggests that the α5SNP results in a partial loss of function in nAChRs when co-assembled [68–70], but genotype-dependent differences in the electrophysiological properties were not observed between WT and α5SNP rats.

It has been demonstrated that the functional expression of the α5 subunit mediates a developmental retraction of apical dendrites in L6 neurons. Neurons in WT mice, but not α5KO mice, exhibit a notable shift towards shorter apical dendrites by early adulthood [52]. Here, we conducted a comparative analysis of the dendritic morphology of L6 RS neurons among the three genotypes using young adult rats. Our findings indicate that the expression of the rs16969968 SNP induces changes in morphological features resembling those observed in α5KO rats, with the majority of neurons having a long apical dendrite that extending into superficial layers. Beyond impacting dendritic morphology, nicotine exposure or nAChR dysfunction have powerful effects on dendritic spine density [37, 53–56]. Consistent with this idea, we observed a gradual decrease in dendritic spine density among L6 RS neurons in WT, α5SNP-carrying, and α5KO rats. The aberrant dendritic morphology and altered spine density of L6 RS neurons in mPFC of α5SNP rats serve as structural correlates of abnormal synaptic plasticity and cortical output resulting from α5 nAChR subunit dysfunction. These changes could potentially contribute to the pathophysiology of neuropsychiatric disorders such as attention-deficit disorder and SZ, which result from the impairment of PFC function [11, 71].

### Potential therapeutic strategies to address cognitive deficits by targeting the α5 nicotinic subunit

It has been shown that in humans the presence of α5SNP decreases resting state functional connectivity of a dorsal anterior cingulate-ventral striatum/extended amygdala circuit. The activity of this circuits predicts addiction severity in smokers and is further impaired in people with mental illnesses [72]. Similarly, in the PFC of α5SNP-expressing mice, lower activity of vasoactive intestinal polypeptide (VIP) interneurons resulted in an increased somatostatin (SOM) interneuron inhibitory drive over L2/3 pyramidal neurons. The decreased activity observed in α5SNP-expressing mice can be reversed by chronic nicotine administration [38]. These studies provide a physiological basis for the tendency of SZ patients carrying α5SNP to self-medicate by smoking. Given the prevalence of the rs16969968 in the general human population, homozygous carriers may benefit from medication that could potentially restore a partial loss of function of the corresponding nAChR subtype. This may help address nicotine addiction and cognitive impairment among patients with SZ. Therefore, it is of great interest to develop pharmacological interventions to ameliorate the relevant dysfunction. Small molecules, such as galantamine, can act as PAMs of channel function in the high-affinity α5*nAChRs. At low concentrations, galantamine binds allosterically to nAChRs and enhances their function; at high concentrations, it also acts as a weak acetylcholinesterase (AChE) inhibitor [23, 50]. Here, we used transgenic rats to compare different genotypes with respect to their nicotinic responses and modulation by galantamine. Following application of 10 µM ACh, only L6 RS neurons in WT rats showed a membrane potential depolarization while neurons in α5SNP and α5KO rats displayed no response. This suggests an essential role of α5*nAChRs in tonic cholinergic neuromodulation mediated by volume transmission [8, 73]. Notably, the application of 1 µM galantamine successfully restored nicotinic responses in L6 neurons of α5SNP rats to WT level. This potentiation effect, however, was not observed in α5KO rats. Our results indicate that galantamine or its derivatives could be a potential pharmacological therapy with high specificity for improving the function of high-affinity nAChRs in populations carrying the rs16969968 polymorphism.

As an AChE inhibitor, galantamine has long been used to treat the cognitive impairments in Alzheimer’s disease [74, 75]. There is growing evidence that galantamine can improve cognitive function in psychiatric disorders including SZ [76–78], major depression [79–81], bipolar disorder [82, 83], and alcohol dependence [84]. These therapeutic effects may result from the allosterically potentiating role of galantamine, which contributes not only to increased nAChR signaling, but also to the enhancement of the release of other neurotransmitters such as glutamate, dopamine and noradrenaline [85–87]. Using the α5SNP rats, we demonstrated that galantamine can correct abnormal nAChR responses in the PFC L6 neuronal circuitry. Further research is needed to explore the corresponding behavioral outcomes. Compared to transgenic mice, transgenic rats emerge as a more suitable animal model for complex behavioral tests. They are an excellent transgenic model for evaluating small molecules designed for the treatment of psychiatric disorders.

### Functional distribution of the α5 subunit is cell type-specific

Two main pyramidal cell populations exist in neocortical layer 6: RS CT neurons and BS CC neurons [46, 47, 49, 61]. By forming reciprocal interactions between the PFC and thalamic nuclei, CT cells are an essential part of the cortico-thalamo-cortical feedback loop, which plays a critical role in cognition [88, 89]. On the other hand, CC neurons send long-range efferents that can innervate other subregions of the PFC, contralateral PFC and other intratelencephalic targets [90, 91]. Previous studies have shown that nicotinic currents in L6 neocortical BS neurons are weaker compared to RS neurons [19, 50]. Here, we provide clear evidence that there is a selective functional expression of *Chrna5* in L6 of mPFC. The small nAChR-mediated depolarization in BS neurons can be attributed to both a low density of α4ß2* nAChRs and the absence of co-assembly with the α5 subunit. Deletion of the α5 subunit resulted in a reduced nicotinic response in RS neurons but not BS neurons. This suggests the absence of α4ß2α5 nAChRs in BS neurons. This finding is consistent with a study that used *in situ* hybridization to demonstrate selective expression of *Chrna5* in CT neurons in mouse primary somatosensory cortex [51]. In α5KO rats, we also observed a weaker nAChR-mediated depolarization in BS neurons compared to RS neurons, indicating a lower density of nAChRs composed solely of α4 and ß2 nAChR subunits.

Recent studies have combined single cell-electrophysiology, morphology and transcriptomics to perform a more comprehensive cell type classification of human cortical neurons [92–95]. However, there is still a lack of studies exploring the combination of transcriptomic and morpho-electric properties in human cortical layer 6, which remains a missing domain. Using spared human cortical samples obtained during brain surgery, we identified the cell type-specific expression of α5*nAChRs in human neocortical layer 6. The nAChR-mediated effects were found to be closely related to both neuronal firing pattern and dendritic morphology. In contrast to L6 RS neurons, which uniformly display upright-oriented apical dendrites, BS neurons exhibit a more heterogeneous morphology including small pyramids, multipolar neurons, and bipolar neurons, consistent with previous studies for CC neurons [96, 97].

In summary, we have shown that L6 RS CT neurons are more efficiently modulated by ACh than BS CC neurons due to the presence of α5*nAChR. As an integral part of the cortico-thalamo-cortical pathway, L6 CT neurons in PFC provide feedback control of thalamo-frontal circuitry, contributing to the maintenance of persistent activity in the higher-order thalamus during behavior [89]. Dysfunctions in the thalamic counterparts of PFC, or thalamo-frontal circuitry have been implicated in various neuropsychiatric disorders including SZ [98–100]. Our findings improve our understanding of cholinergic modulation of neuronal microcircuits involved in cognition and highlight a potential pharmacological target for restoring nicotinic signaling under pathological conditions.

## Author contributions

D.F., D.Y., and G.Q designed the experiments. D.Y. and G.Q. carried out the patch-clamp recording experiments from human and rat slices and performed electrophysiological data analysis. D.Y. performed morphological and spine analysis. D.D performed surgeries on human patients. U.M. provided the transgenetic rat models. D.Y. and D.F. wrote the manuscript with the inputs from all authors. All authors have given approval for the final version of the manuscript.

## Acknowledgement

We would like to thank Werner Hucko for excellent technical assistance. We thank Dr. Karlijn van Aerde for custom-written macros in Igor Pro software. We thank Yutong Wu for reconstructing 3D dendritic morphologies of both rat and human neurons. We thanks Dr. Henner Koch and Aniella Bak for kind help with human slice preparations. We are grateful for funding support from the European Union’s Horizon 2020 Framework Programme for Research and Innovation under the Framework Partnership Agreement No. 650003 (HBP FPA) to D.F.

