## Supplementary Materials for "Linking Altered Neuronal and Synaptic Properties to Nicotinic Receptor Alpha5 Subunit Gene Dysfunction: A Translational Investigation in Rat mPFC and Human Cortical Layer 6"

### These authors share the first authorship

**Tab. S1 Electrophysiological and morphological properties of L6 RS neurons in WT, α5SNP and α5KO rats.**

Italic bold font indicates significant differences; *P < 0.05, **P < 0.01, ***P < 0.001 for Wilcoxon Mann-Whitney U test.

| | WT | SNP | KO | Mann-Whitney Test | | |
|---|---|---|---|---|---|---|
| *Electrophysiological properties* | *n = 13* | *n = 10* | *n = 19* | *WT vs. α5SNP* | *WT vs. α5KO* | *α5SNP vs. α5KO* |
| Resting membrane potential (mV) | -71.7 ± 4.2 | -69.7 ± 4.0 | -65.0 ± 7.0 | 0.4833 | **0.0035 | 0.0771 |
| Rheobase current (pA) | 87.7 ± 24.9 | 86.0 ± 29.1 | 58.4 ± 29.3 | 0.7973 | **0.0090 | *0.0250 |
| Input resistance (MΩ) | 251 ± 44 | 255 ± 70 | 330 ± 60 | 0.6926 | ***7.7E-05 | **0.0024 |
| Voltage sag (mV) | 0.70 ± 0.69 | 0.65 ± 0.34 | 1.65 ± 1.82 | 0.5538 | *0.0491 | 0.08289 |
| Time constant (ms) | 32.0 ± 11.1 | 29.5 ± 5.0 | 32.7 ± 8.5 | 0.7844 | 0.6500 | 0.4357 |
| AP half- width (ms) | 1.00 ± 0.13 | 1.08 ± 0.12 | 1.06 ± 0.11 | 0.1259 | 0.0859 | 0.5492 |
| AP amplitude (mV) | 94.5 ± 5.2 | 92.9 ± 8.0 | 89.8 ± 6.4 | 0.7381 | *0.0302 | 0.5125 |
| AP threshold (mV) | -35.2 ± 7.5 | -35.4 ± 5.8 | -34.0 ± 6.1 | 0.9758 | 0.4315 | 0.6038 |
| AHP amplitude (mV) | 16.4 ± 5.4 | 15.9 ± 1.9 | 18.0 ± 3.8 | 0.8315 | 0.0916 | 0.0559 |
| AP latency (ms) | 219 ± 173 | 344 ± 145 | 339 ± 129 | 0.0875 | *0.0177 | 0.8747 |
| Adaptation ratio ($ISI_2$/ $ISI_{10}$) | 0.87 ± 0.2 | 0.91 ± 0.12 | 0.94 ± 0.14 | 0.5006 | 0.4652 | 0.7261 |
| Frequency- current slope (Hz/100 pA) | 14.9 ± 4.6 | 12.4 ± 2.7 | 16.2 ± 5.2 | 0.2692 | 0.4770 | *0.0406 |
| *Morphological properties* | *n = 10* | *n = 8* | *n = 10* | *WT vs. α5SNP* | *WT vs. α5KO* | *α5SNP vs. α5KO* |
| Somatic area (µm$^2$) | 182 ± 52 | 194 ± 45 | 178 ± 42 | 0.3599 | 0.9705 | 0.3154 |
| Length of apical dendrite (mm) | 3.5 ± 1.0 | 4.0 ± 1.1 | 3.5 ± 0.6 | 0.4082 | 0.6305 | 0.3599 |
| Total length of basal dendrites (mm) | 1.4 ± 0.5 | 2.0 ± 0.5 | 1.5 ± 0.3 | *0.0266 | 0.2475 | *0.0343 |
| Mean length of basal dendrite (mm) | 0.24 ± 0.05 | 0.27 ± 0.08 | 0.20 ± 0.04 | 0.4598 | 0.1655 | *0.0434 |
| No. of basal dendrites | 5.8 ± 1.5 | 7.5 ± 1.3 | 7.3 ± 1.1 | *0.0290 | *0.0214 | 0.8287 |
| H-fieldspan of dendrite (mm) | 0.40 ± 0.12 | 0.37 ± 0.05 | 0.31 ± 0.07 | 0.4997 | *0.0371 | 0.1051 |
| V-fieldspan of dendrite (mm) | 0.81 ± 0.27 | 0.97 ± 0.15 | 1.00 ± 0.09 | 0.2643 | 0.1595 | 0.9829 |
| Aspect ratio of dendritic fieldspan | 2.2 ± 0.9 | 2.7 ± 0.5 | 3.4 ± 0.9 | 0.2031 | *0.0147 | 0.2031 |

**Table S2. Patient demographic and clinical data for the tissue used in this study.**

| Patient Number | Patient Age | Diagnosis | Gender | Brain region |
|---|---|---|---|---|
| **1** | 34 | Ganglioglom | Male | Frontal |
| **2** | 66 | Glioblastoma multiforme | Male | Temporal |
| **3** | 37 | Hippocampus Sclerosis | Female | Temporal |
| **4** | 23 | Normal | Male | Parietal |
| **5** | 32 | Hippocampus Sclerosis | Male | Temporal |
| **6** | 34 | Hippocampus Sclerosis | Male | Temporal |

**Tab. S3 Comparison of electrophysiological and morphological properties between human L6 RS and BS neurons.**

Italic bold font indicates significant differences; *P < 0.05, **P < 0.01, ***P < 0.001 for Wilcoxon Mann-Whitney U test.

| | RS neurons | BS neurons | Mann-Whitney Test |
|---|---|---|---|
| *Electrophysiological properties* | ***n = 11*** | ***n = 11*** | |
| **Resting membrane potential (mV)** | -70.6 ± 5.5 | -72.6 ± 5.5 | 0.8470 |
| **Rheobase current (pA)** | 72.7 ± 44.3 | 81.8 ± 28.9 | 0.8428 |
| **Input resistance (MΩ)** | 232.5 ± 112.0 | 184.5 ± 64.1 | 0.4009 |
| **Voltage sag (mV)** | 1.7 ± 1.1 | 0.9 ± 0.5 | ****0.0281*** |
| **Time constant (ms)** | 42.9 ± 16.7 | 38.0 ± 10.2 | 0.7477 |
| **AP half- width (ms)** | 1.09 ± 0.21 | 1.02 ± 0.17 | 0.4675 |
| **First AP amplitude (mV)** | 92.5 ± 8.0 | 94.8 ± 4.8 | 0.2816 |
| **Second AP amplitude (mV)** | 90.7 ± 7.5 | 79.0 ± 6.1 | ******0.0006*** |
| **Third AP amplitude (mV)** | 90.1 ± 6.9 | 86.7 ± 4.9 | 0.2512 |
| **AP threshold (mV)** | -43.2 ± 2.8 | -39.9 ± 5.4 | 0.0879 |
| **AHP amplitude (mV)** | 16.9 ± 4.1 | 13.7 ± 4.2 | 0.0879 |
| **AP latency (ms)** | 223.9 ± 84.6 | 265.3 ± 61.0 | 0.1164 |
| **Adaptation ratio ($ISI_2/ISI_{10}$)** | 0.71 ± 0.16 | 0.31 ± 0.15 | ******3.7E-05*** |
| **Adaptation ratio ($ISI_3/ISI_{10}$)** | 0.90 ± 0.11 | 1.33 ± 0.43 | *****0.0022*** |
| **Frequency- current slope (Hz/100 pA)** | 13.5 ± 5.3 | 5.8 ± 1.2 | ******4.0E-05*** |
| *Morphological properties* | ***n = 9*** | ***n = 9*** | |
| **Somatic area (µm²)** | 356 ± 87 | 321 ± 109 | 0.3401 |
| **Length of apical dendrite (mm)** | 6.7 ± 3.6 | 5.9 ± 3.0 | 0.6048 |
| **Total length of basal dendrites (mm)** | 5.7 ± 2.0 | 6.2 ± 3.0 | 0.7962 |
| **length of longest basal dendrite (mm)** | 1.6 ± 0.8 | 4.0 ± 2.6 | ****0.0498*** |
| **No. of basal dendrites** | 5.7 ± 1.0 | 7.0 ± 2.8 | 0.3009 |
| **H-fieldspan of dendrite (mm)** | 0.55 ± 0.14 | 0.81 ± 0.32 | 0.0770 |
| **V-fieldspan of dendrite (mm)** | 1.57 ± 0.33 | 1.18 ± 0.37 | ****0.0315*** |
| **Aspect ratio of dendritic fieldspan** | 2.9 ± 0.6 | 1.5 ± 0.5 | ******0.0005*** |
